# Individual bat viromes reveal the co-infection, spillover and emergence risk of potential zoonotic viruses

**DOI:** 10.1101/2022.11.23.517609

**Authors:** Jing Wang, Yuan-fei Pan, Li-fen Yang, Wei-hong Yang, Chu-ming Luo, Juan Wang, Guo-peng Kuang, Wei-chen Wu, Qin-yu Gou, Gen-yang Xin, Bo Li, Huan-le Luo, Yao-qing Chen, Yue-long Shu, Deyin Guo, Zi-Hou Gao, Guodong Liang, Jun Li, Edward C. Holmes, Yun Feng, Mang Shi

## Abstract

Bats are reservoir hosts for many zoonotic viruses. Despite this, relatively little is known about the diversity and abundance of viruses within bats at the level of individual animals, and hence the frequency of virus co-infection and inter-species transmission. Using an unbiased meta-transcriptomics approach we characterised the mammalian associated viruses present in 149 individual bats sampled from Yunnan province, China. This revealed a high frequency of virus co-infection and species spillover among the animals studied, with 12 viruses shared among different bat species, which in turn facilitates virus recombination and reassortment. Of note, we identified five viral species that are likely to be pathogenic to humans or livestock, including a novel recombinant SARS-like coronavirus that is closely related to both SARS-CoV-2 and SARS-CoV, with only five amino acid differences between its receptor-binding domain sequence and that of the earliest sequences of SARS-CoV-2. Functional analysis predicts that this recombinant coronavirus can utilize the human ACE2 receptor such that it is likely to be of high zoonotic risk. Our study highlights the common occurrence of inter-species transmission and co-infection of bat viruses, as well as their implications for virus emergence.

## INTRODUCTION

Bats (order Chiroptera) are hosts for a larger number of virus species than most mammalian orders^1^, and are the natural reservoirs for several emerging viruses that cause infectious disease in human^2^. Recently, there has been considerable research effort directed toward exploring viral diversity in bats as a means to identifying potential zoonotic infections^3^. These studies have greatly expanded the diversity of known bat-borne viruses and identified an array of potentially emerging viruses. However, despite the growing body of work on bat viruses, little is known about the underlying drivers of virus diversity within these animals, nor of the extent patterns of viral co-infection and the frequency of viral spillover among bat species^4,5^.

Current virus discovery studies typically pool individual bats by species or by sampling location (e.g., ref ^6,7^). Although of great utility, this hinders mechanistic insights due to insufficient resolution. As such, studying the bat virome at the scale of individual animals can help us better understand the diversity and emergence of bat-borne viruses^4^. For example, the co-infection of phylogenetically related viruses within an individual host facilitates the occurrence of recombination^8^, which may in turn have contributed to the emergence of a number of zoonotic viruses (e.g. SARS-CoV^9^). Importantly, the frequency of virus co-infection in bats can be resolved through the study of the viromes of individual animals. Resolution at the scale of individual animals is also required to better understand the frequency and determinants of virus spillover among bats^4,10^. For example, correlating host traits at the individual level with the probability of cross-species transmission is an important way to reduce confounding effects.

Many previous studies of bat viruses have preferentially targeted relatives of known human pathogens^3^. Although time and cost-effective, this necessarily limits our ability to discover novel zoonotic viruses. In contrast, other studies have utilized metagenomics approaches to explore the total bat virome (e.g., ref ^7^), with meta-transcriptomic sequencing demonstrating great utility as a means to characterise the total diversity of viruses without *a priori* knowledge of which viruses are present^11,12^.

Yunnan province in southwestern China has been identified as a hotspot for the diversity of bat species and bat-borne viruses. A number of highly pathogenic viruses have been detected there, including close relatives of SARS-CoV-2, such as bat viruses RaTG13^13^, RpYN06^14^, and RmYN02^15^, as well as relatives of SARS-CoV, such as WIV1^16^ and Rs4231^17^. It has been hypothesized that the presence of mixed roosts of bats in Yunnan (i.e., multiple bat species occupying the same roost) contributes to the frequent cross-species transmission of viruses, promoting their recombination and ultimately leading to transient spillovers or successful cross-species transmissions^17^. Thus, wild bat populations in Yunnan provide a unique opportunity to study the diversity, spillover and emergence risk of bat-borne viruses.

We performed intensive field sampling of individual wild bats in Yunnan province, China. In particular, we characterised the total mammal-associated virome of wild bats at the scale of individual animals using unbiased meta-transcriptomic sequencing. We then explored the cross-species transmission of viruses among individual animals from different species and quantitatively tested how host phylogeny and geographic (i.e. sampling) location may impact the probability of cross-species transmission. Finally, we identified viruses of potentially high emergence risk and evaluated their pathogenic potential using a combination of phylogenetic analysis and *in sillico* molecular dynamics simulations.

## RESULTS

### Characterization of the bat viromes

Between 2015 and 2019, rectum samples were collected from 149 individuals of bat in six counties/cities, Yunnan province, China, representing six genera and 15 species (Fig. 1; Table S1). Total RNA was extracted and sequenced individually for each individual bat. Meta-transcriptomic sequencing yielded an average of 41,789,834 clean non-rRNA reads for each animal. In total, 1,048,576 contigs were *de novo* assembled from non-rRNA reads, among which the 36,029 contigs annotated as viruses were used for virome characterization.

**Fig. 1.**
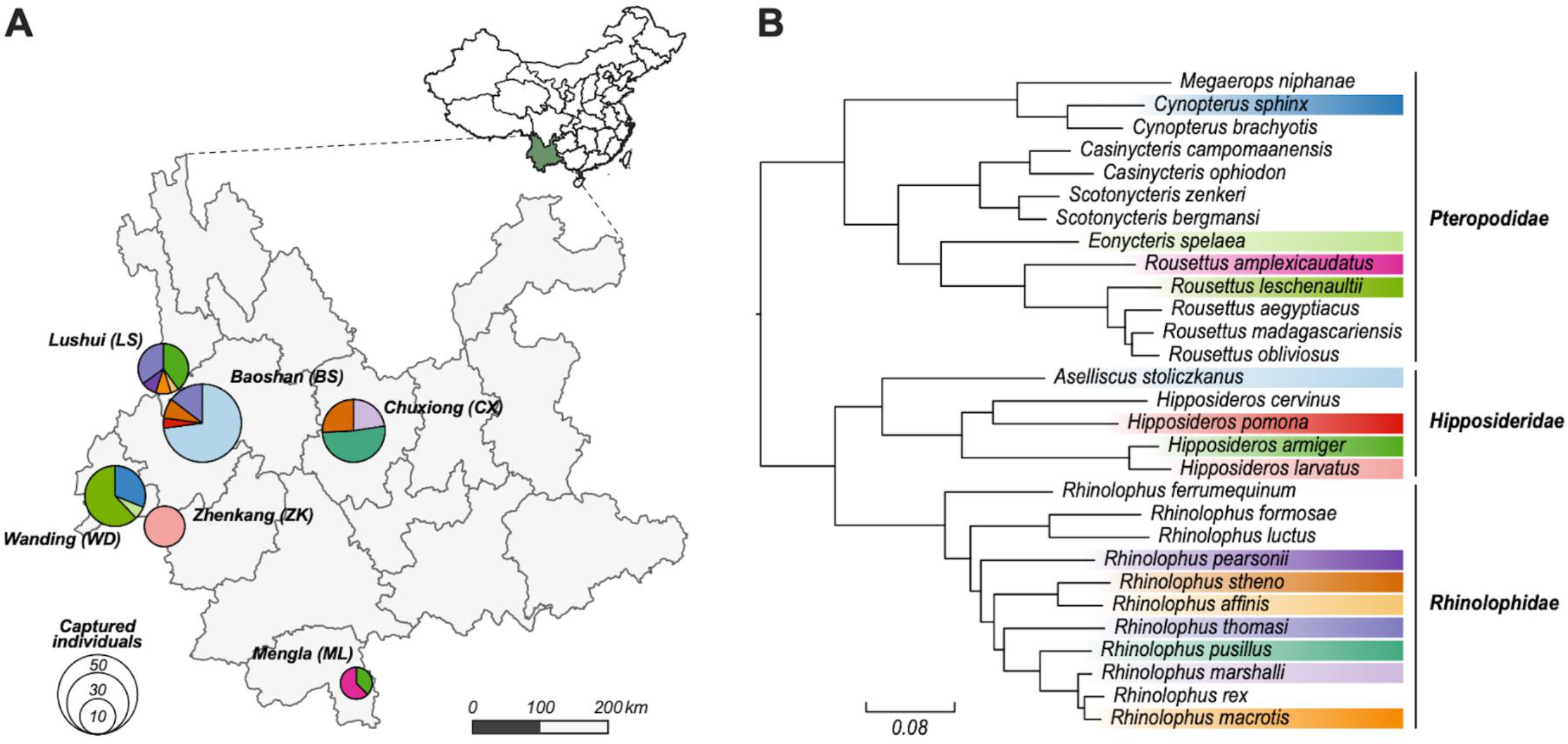
Overview of the samples analysed in this study. (**A**) Locations in Yunnan province China where bat samples were taken. Pie charts indicate the composition of bat species sampled at each location, while the total area of the pies are proportional to number of captured individuals. Colours indicate different bat species, which are consistent with the colouring scheme in plot B. (**B**) Phylogeny of bats, including those sampled as part of this study. The tree was estimated using nucleotide sequences of bat COI gene utilising a maximum likelihood (ML) method. Coloured strips indicate the bat species sampled in this study.

We focused on characterizing the mammal-associated viromes of bats (Fig. 2), which include the RNA and DNA viral families or genera that are known to infect mammalian hosts (rather than those viruses more likely associated with bat diet or microbiome). In total, we identified 41 mammal-associated virus species belonging to 11 families (Table S2). Most of the viruses detected were RNA viruses, comprising 32 of the 41 viral species. The *Reoviridae* was the most prevalent viral family, present in 27.5% of individuals sampled, followed by the *Picornaviridae* (12.1%) and the *Coronaviridae* (8.7%) (Fig. 3C). The prevalence of the remaining viral families was relatively low (≤ 4%).

**Fig. 2.**
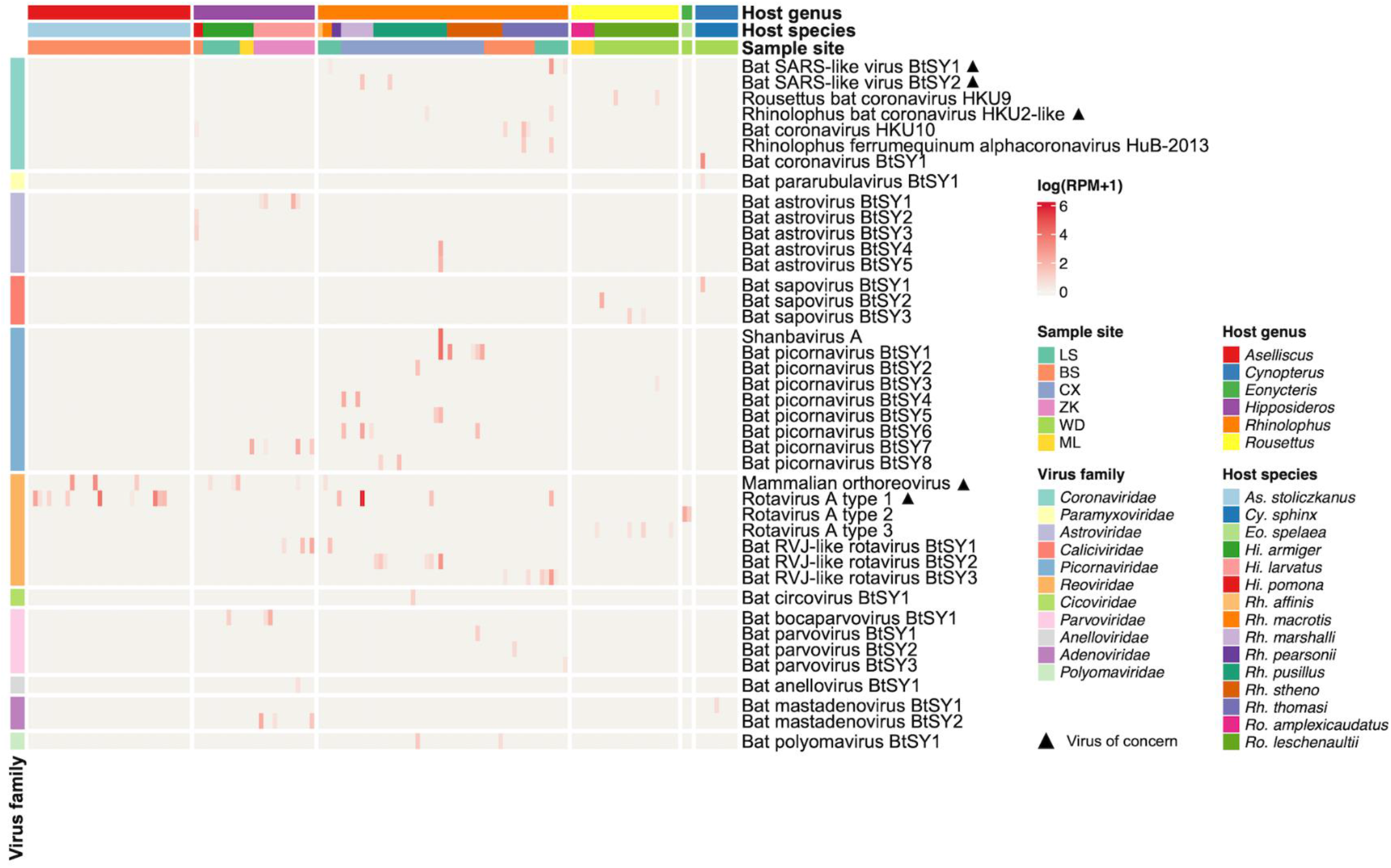
Characterization of the mammal-associated virome of bats. The heatmap displays the distribution and abundance of mammal-associated viruses in individual bats. Each column represents an individual bat, while each row represents a virus species. The abundance of viruses in each individual is represented as a logarithm of the number of mapped reads per million total reads (RPM). Sampling site, host taxonomy (species and genus) and virus taxonomy are shown as coloured strips at top and left, respectively. Black triangle marks indicate “viruses of concern”, defined as those that are closely related to known human or livestock pathogens (>90% amino acid similarity in RNA-dependent RNA polymerase).

**Fig. 3.**
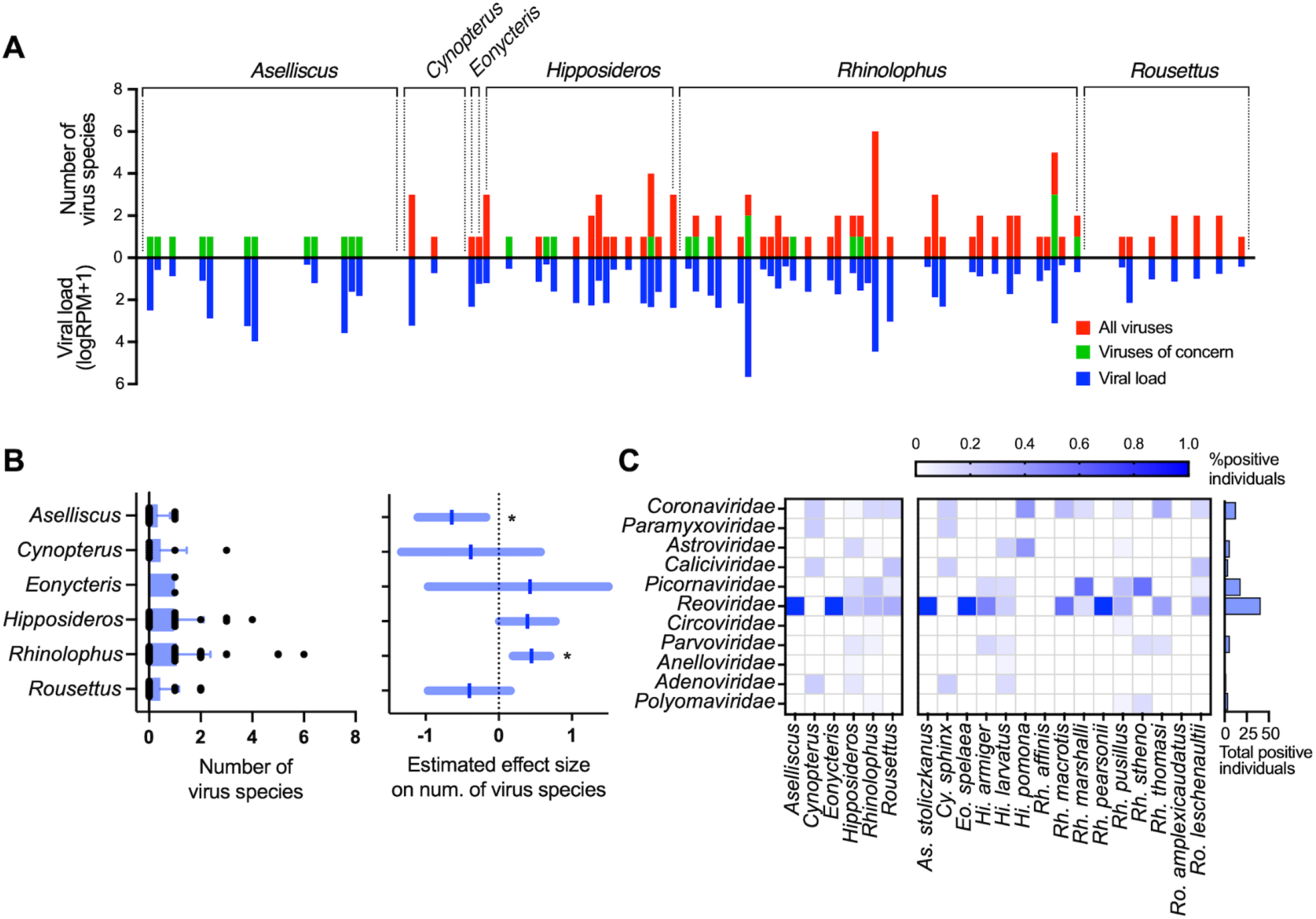
Comparison of mammal-associated virus diversity among different bat taxa. (**A**) Virus load and the number of virus species in individual bats. Red bars, the total number of mammal-associated viruses per host. Green bars, number of viruses of concern per host. Blue bars represent viral load per host as logarithm of the sum of total viral RPM. (**B**) Left, comparison of the number of viruses per individual host among six bat genera (mean+SD). Right, estimated effect size of each bat genus on the number of virus species per individual bat by Poisson regression (mean±95%CI). Stars indicate significant effects (in which zero is not included in the 95%CI). (**C**) Comparison of the prevalence of 11 viral families among different host genera (left block) and species (right block).

We next quantified the virus load and number of virus species for each individual bat (Fig. 3). Of the 149 individual bats analysed, 70 were positive for at least one virus species (positive rate 47.0%). We consider those with relatively high viral load (reads per million total reads > 1) as true positives. Among positive individuals, one-third were infected by more than one viral species (1.5 viral species per individual on average). The number of virus species per individual was uneven among host genera (Fig. 3B). We used Poisson regression to estimate the effect of host genera on the number of virus species. This revealed that *Rhinolophus* bats carried significantly more virus species per individual than average, whereas *Aselliscus* bats carried significantly less.

### Cross-species transmission of viruses among bats

We identified 12 virus species that are shared among different bat species, accounting for the possibility of index-hopping (Fig. 4A). The 12 species identified belong to the *Coronaviridae* (4 species), *Reoviridae* (3), *Picornaviridae* (3), *Parvoviridae* (1) and *Polyomaviridae* (1). Rotavirus A type 1 (RVA1) and Mammalian orthoreovirus (MRV) had the broadest host range, being detected in five and four bat species, respectively, with both these viruses shared among the bat families Hipposideridae and Rhinolophidae. The remaining 10 viral species were only found in two bat species, and most were only shared among animals from the same host genus, with the exception of bat coronavirus HKU10 (BtCoV-HKU10, present in *Hi. pomona* and *Rh. thomasi*) and Bat RVJ-like rotavirus BtSY1 (BtRV1, present in *Hi. larvatus* and *Rh. macrotis*).

**Fig. 4.**
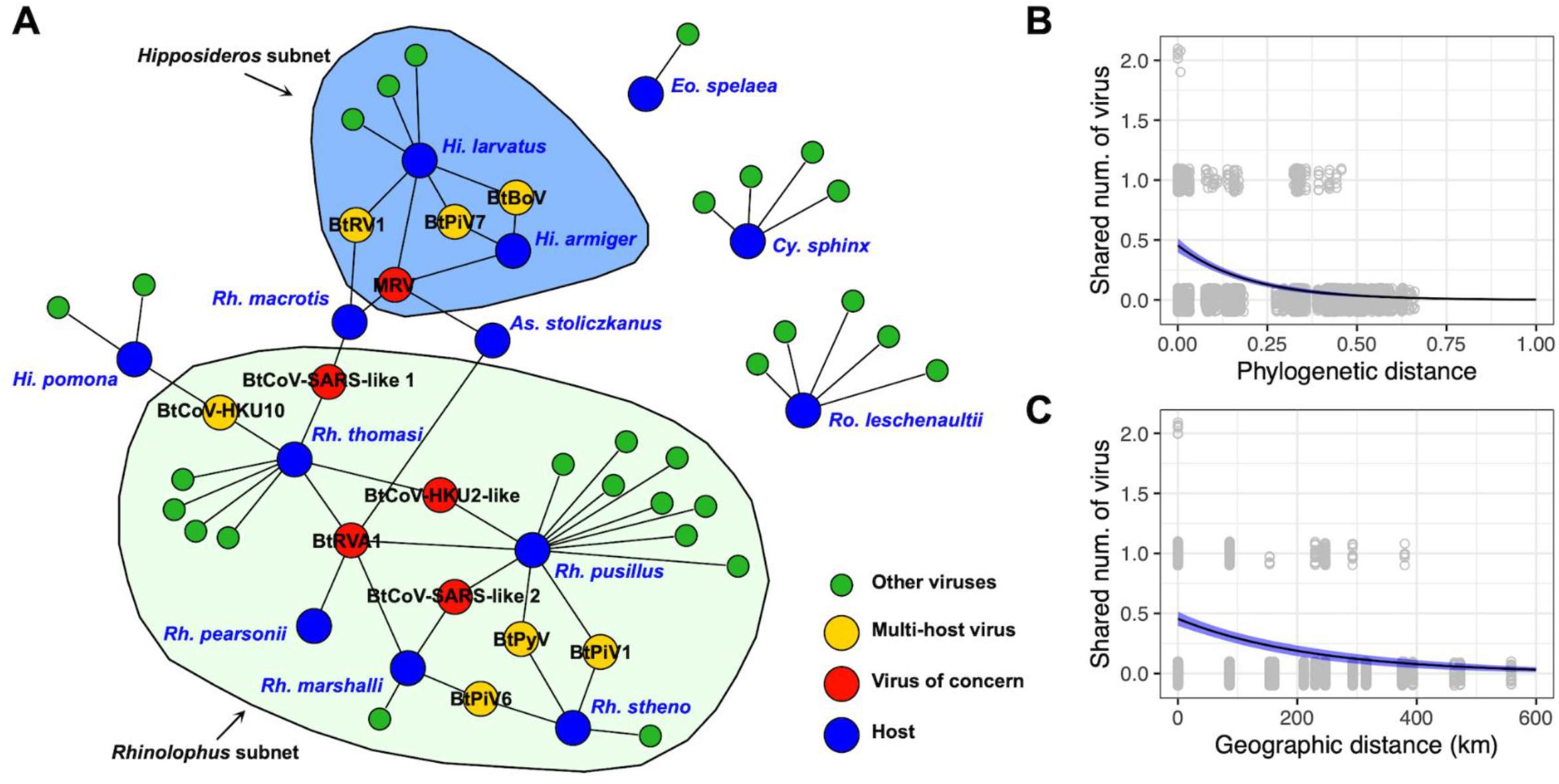
The virus-sharing network of bats. (**A**) The virus-sharing network reveals connectivity among viromes of different bat taxa. Viruses of concern and putative cross-species transmissions are shown in different colours. Two network modules (subnets) were detected with a network betweenness-based criterion and are visualised by coloured areas. (**B, C**) The relationship between the number of shared viruses with phylogenetic (B) or geographic distance (C) between pairs of host individuals. Phylogenetic distance is calculated as the sum of phylogenetic tree branch length between a pair of hosts, and the tree was estimated with nucleotide sequences of the COI gene employing a maximum likelihood method. The line and blue area is the estimated partial effect and standard error of phylogenetic or geographic distance by Poisson regression.

The number of viral species found in two bat species ranged from zero to three. Partial Mantel tests showed that more closely phylogenetically related or closely geographically located bat individuals had more similar virome compositions and had more virus species in common (Table S4). For example, the viromes of *Rhinolophus* or *Hipposideros* bats form two network modules, in which individuals within the same genus are more inter-connected (i.e., shared more viruses) than individuals from different genera (Fig. 4A). A Poisson regression analysis showed that the number of virus species shared between pairs of bats was significantly associated with both the phylogenetic and the geographic distance of hosts, after controlling for the confounding effect of date of sampling (Fig. 4B, C).

### Phylogenetic analysis identifies viruses of potentially high emergence risk

Phylogenetic analysis identified five viral species what were closely related to known human or livestock pathogens, which we termed “viruses of concern” (Fig. 5 and Table S5). The five viruses of concern belong to two viral families – the *Coronaviridae* (three species) and the *Reoviridae* (two species). Notably, all the five viruses were detected in more than one bat species, and their prevalence was relatively high, especially Mammalian orthoreovirus and Rotavirus A type 1 (Table S5).

**Fig. 5.**
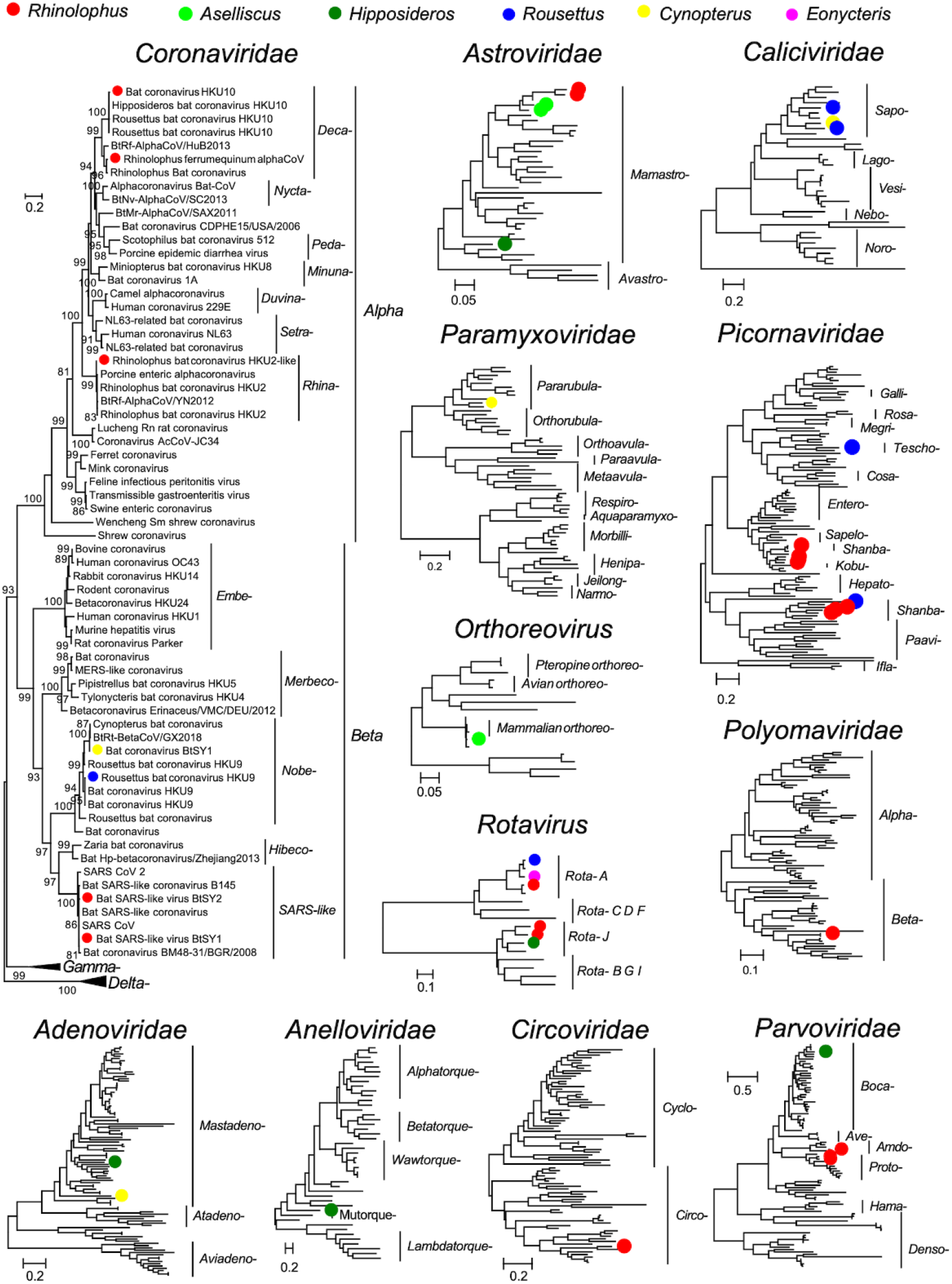
Evolutionary relationships of 11 viral families detected in our study. Phylogenetic trees were estimated using a maximum likelihood method based on conserved replicase protein (RNA viruses: RdRp, *Polyomaviridae*: LTAg, *Anelloviridae*: ORF1 protein, *Parvoviridae*: NS1, and other DNA viruses: DNA pol). Trees were midpoint rooted, and the branch length indicates number of nucleotide substitutions per site. Dots indicate viruses detected in our samples, and colours represent host genus.

The three coronaviruses were closely related to highly pathogenic viruses that infect humans or swine. A phylogenetic analysis using the conserved RdRp protein revealed that both Bat SARS-like virus BtSY1 and BtSY2 belong to the subgenus *Sarbecovirus* of betacoronaviruses and are closely related to human SARS-CoV (>90% nucleotide identity). Notably, other key functional genes (e.g., NTD, RBD, N) of Bat SARS-like virus BtSY2 were more closely related to SARS-CoV-2 (i.e., the early Wuhan-Hu-1 reference strain), indicative of a past history of recombination. We present further analysis of the evolutionary history and zoonotic potential of this virus below. The other coronavirus, which was Rhinolophus bat coronavirus HKU2-like, belonged to the genus *Alphacoronavirus* and was closely related to SADS-CoV of pigs in the RdRp gene.

The remaining two viruses of concern from the *Reoviridae* are known species -Mammalian orthoreovirus and Rotavirus A type 1. In total, three different types of Rotavirus A were identified according to the RdRp phylogeny. The nucleotide identity between their RdRp was less than 80%, so we demarcated them as three different types. Of these, Rotavirus A type 1 was most closely related to human Rotavirus A, while Rotavirus Type 2 and 3 belong to viral lineages associated with *Eonycteris* and *Rousettus* bats, respectively (Fig. S1).

As well as viruses of concern, 28 viral species can be classed as newly discovered viruses of mammals. The *Picornaviridae* (n = 8) contained the highest number of the newly discovered viral species, followed by the *Astroviridae* (n = 5), *Parvoviridae* (n = 4), and *Caliciviridae* (n = 3).

### The evolutionary history and zoonotic potential of two SARS-related coronavirus

We next evaluated the evolutionary history and zoonotic potential of the two SARS-related coronaviruses detected in our samples: Bat SARS-like virus BtSY1 and BtSY2 (for simplicity referred to as BtSY1 and BtSY2, respectively, in the following text) (Fig. 6). Phylogenetic trees were estimated using the nucleotide sequences of key genes: the RdRp (RNA-dependent RNA polymerase), N-terminal domain (NTD) and receptor-binding domain (RBD) of spike protein, and the nucleoprotein (N). This analysis revealed that in the NTD, RBD and N gene trees, BtSY1 clustered with SARS-CoV forming an “S-1” clade, while BtSY2 clustered with SARS-CoV-2 forming the “S-2” clade. Notably, while BtSY1 remained in the S1 clade in the phylogeny of the RdRp gene, BtSY2 also fell into the S-1 clade. Hence, BtSY2 appears to be a recombinant between the S-1 and S-2 lineages.

**Fig. 6.**
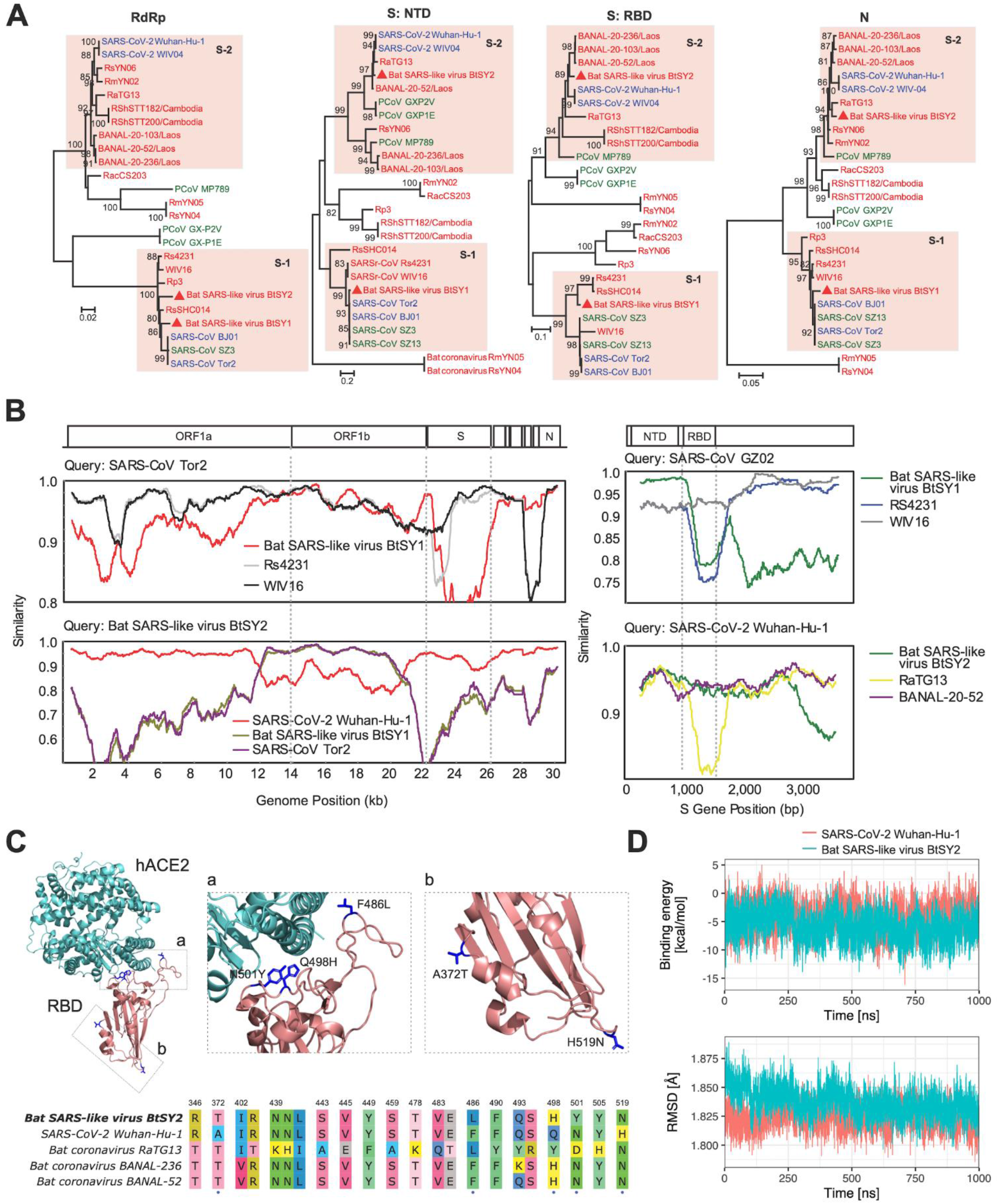
Phylogenetic and structural analysis of a potentially zoonotic SARS-related coronavirus detected in our samples. (**A**) Phylogenetic trees of four key functional genes of SARS-related coronaviruses. Colours of virus strain names indicate the host taxa where the viruses were detected. Red: bats, blue: human, green: others. (**B**) Recombination analysis of SARS-related coronaviruses at the whole genome and spike protein scales. (**C**) Top, homology-modelling structure of the receptor-binding domain (RBD) of Bat SARS-like virus BtSY2 in complex with human angiotensin-converting enzyme 2 (hACE2). Blue-coloured residues on RBD indicate amino-acid substitutions compared with SARS-CoV-2 Wuhan-Hu-1. Bottom, alignment of RBD sequences (residues T333 to G526 of spike protein) of Bat SARS-like virus BtSY2, SARS-CoV-2 Wuhan-Hu-1 and two closely related bat coronavirus. Only polymorphic sites are shown. The five amino acid differences in the RBD of Bat SARS-like virus BtSY2 compared with SARS-CoV-2 Wuhan-Hu-1 are marked with blue dots. (**D**) Molecular dynamics simulation results of stability (top) and binding energy (bottom) of Bat SARS-like virus BtSY2 RBD-hACE2 complex. BtSY1 and BtSY2 are abbreviations for Bat SARS-like virus BtSY1 and BtSY2, respectively.

At the scale of the whole genome, BtSY1 generally exhibited the highest genetic identity to human SARS-CoV viruses (93%). Indeed, in comparisons to previously identified SARS-related viruses (i.e., WIV16, Rs4231), BtSY1 shared highest identity with human SARS-CoV viruses in ORF1b (nsp13 and nsp15) and the NTD, although it was relatively more distant in ORF1a and the RBD, as well as in the S2 region of S gene (Fig. 6B). Specifically, it exhibited 98.13% similarity with SARS-CoV in the NTD, but only 88.61% identity in the RBD domain.

In marked contrast, BtSY2 shared 92% genetic identity with SARS-CoV-2 at the whole genome scale, although with the occurrence of recombination. Indeed, we identified potential recombination at positions 12035-20708bp, which encodes ORF1a (nsp7∼nsp11) and ORF1b (nsp12∼nsp14), with this region instead showing strong sequence similarity to SARS-CoV (92.3%). The remainder of its genome is very similar to SARS-CoV-2, particularly in the region encoding the NTD and RBD (95.15% and 93.70%, respectively), although no furin cleavage site was detected in the spike protein (Fig. 6B).

To evaluate the human-receptor-binding potential of BtSY2, we inferred the structure of its RBD using a homology-modelling approach and performed molecular dynamics simulations (Fig. 6). This revealed that there are only five amino-acid substitutions in the RBD in comparison to the SARS-CoV-2 strain Wuhan-Hu-1, with three of these located at the interface of RBD-hACE2 complex (i.e., the receptor-binding motif) (Fig. 6C). Molecular dynamics simulations further revealed that the binding stability and energy of the RBD-hACE2 complex are very similar between BtSY2 and SARS-CoV-2 Wuhan-Hu-1 (Fig. 6D), suggesting that BtSY2 may be able to utilize human ACE2 receptor for cell entry.

## DISCUSSION

We have characterised the mammalian-associated virome of individual bats. This revealed an unexpectedly high frequency of co-infection, with approximately one-third of the virus-positive individuals simultaneously infected by two or more viruses. The frequency of co-infection in individual bats has seldom been investigated, and only a few studies have explored the co-infection of specific viral species using consensus PCR methods (e.g., paramyxovirus, ref ^18^). As such, this study provides the first empirical evidence for co-infection using an unbiased omics approach. Co-infection is prerequisite for virus recombination or reassortment^8^, and the gut microbiome can facilitate the recombination of enteric viruses^19^. Hence, the high frequency of co-infection observed here suggests that recombination and reassortment are very likely to happen within individual bats, which in turn may facilitate the emergence of zoonotic viruses^9^.

Our results also revealed frequent virus spillover among different bat species, identifying 12 different viral species from different families that infect multiple host species. The ability of viruses to jump host species boundaries appears to be a near universal trait among viruses^20^. Our results are of note because they show that the probability of virus spillover among pairs of host individuals is negatively associated with host phylogenetic and geographic distance, thereby supporting the hypothesis that phylogenetically related or spatially closely located hosts share more viruses^21,22^. The frequent virus spillover among phylogenetically related or spatially co-located bats provides an opportunity for viromes of different bat species to exchange, further expanding genetic diversity of circulating viruses.

We identified two SARS-related coronaviruses in *Rhinolophus* bats (*Rh. marshalli, Rh. pusillus Rh. thomasi*, and *Rh. macrotis*) which we suggest are at particular risk for emergence. One of the SARSr-CoV - Bat SARS-like coronavirus BtSY2 (i.e., BtSY2) - is related to both SARS-CoV and SARS-CoV-2 and likely to have a history involving recombination. Notably, there are only five amino acid differences in the receptor-binding domain (RBD) of spike protein of this virus compared with SARS-CoV-2 strain Wuhan-Hu-1^23^, which makes it the closest relative to SARS-CoV-2 found in China in this particular genomic region. In contrast, the nsp7∼nsp11 proteins of ORF1a and nsp12∼nsp14 protein of ORF1b were closely related to SARS-CoV, indicating that these genes were likely to be acquired from another SARSr-CoV. The remainder of its genome was closely related to SARS-CoV-2 and to several bat coronavirus previously found in Yunnan, including RaTG13^13^, RmYN02^15^, and RpYN06^14^ that are all close relatives of SARS-CoV-2. Together, these findings strongly suggest that virus spillover and co-infection in related bat species contribute to the recombination of potentially pathogenic coronavirus and could possibly facilitate virus emergence in other species.

Functional analysis indicated that Bat SARS-like coronavirus BtSY2 likely has the ability to the bind human ACE2 receptor, and even has slightly higher affinity than SARS-CoV-2 Wuhan-Hu-1. Three of the five substitutions in the RBD - Q498H, N501Y and H519N - have been reported to increase affinity to human ACE2^24^, and notably, the N501Y substitution is present in the Alpha, Beta, Gamma and Omicron variants of SARS-CoV-2. In addition, we found that the nsp7-nsp14 proteins (in which nsp12 is the replicase, i.e., RdRp) of BtSY2 were closely related to those of SARS-CoV. A comparative study showed that SARS-CoV can replicate more rapidly than SARS-CoV-2 *in vitro*^25^, while another suggested that nsp14 is likely associated with virulence^26^. Hence, these data tentatively suggest that BtSY2 may be able to replicate rapidly with similar virulence as SARS-CoV. Although this issue merits further consideration, this virus is potentially of high risk of emergence and so should be monitored carefully.

We identified another four viruses of concern, likely to be pathogenic in humans or livestock. Bat SARS-like virus BtSY1 is closely related to SARS-CoV^27,28^. Rhinolophus bat coronavirus HKU2-like is closely related to SADS-CoV, which causes severe diarrhea and death in swine^29,30^, Rotavirus A causes diarrhea in humans^31,32^, while Mammalian orthoreovirus is known to have a broad host range and cause diarrhea in swine^33,34^. Interestingly, all these viruses of concern were found in more than one bat species in our samples, suggesting that these potentially zoonotic viruses may have a broader host range or have a higher rate of spillover than other viruses.

## METHODS

### Ethics Statement

This research, including the procedures and protocols of specimen collection and processing, was reviewed and approved by the Medical Ethics Committee of the Yunnan Institute of Endemic Diseases Control and Prevention. (No. 20160002).

### Sample collection

A total of 149 rectum samples from bats were collected from six counties/cities in Yunnan province, China between 2015 and 2019. Bats were trapped using net traps and were primarily identified according to morphological criteria and confirmed by a barcode gene (COI) in the meta-transcriptomics analysis. The bats collected belonged to 15 species. The majority were from the genus *Rhinolophus* (n=54) and comprised *Rhinolophus pusillus* (n=16), *Rhinolophus thomasi* (n=14), *Rhinolophus stheno* (n=12), *Rhinolophus marshalli* (n=7), *Rhinolophus pearsonii* (n=2), *Rhinolophus macrotis* (n=2), and *Rhinolophus affinis* (n=1). The genus *Hipposideros* (n=26) animals comprised *Hipposideros larvatus* (13), *Hipposideros armiger* (11), and *Hipposideros pomona* (2). The genus *Rousettus* (n=23) animals comprised *Rousettus leschenaultia* (n=18) and *Rousettus amplexicaudatus* (n=5). The *Aselliscus* (n=35), *Cynopterus* (n=9) and *Eonycteris (n=2)* genera animals only contained *Aselliscus stoliczkanus* (n=35), *Cynopterus sphinx* (n=9), *and Eonycteris spelaea* (n=2), respectively. All rectum samples were collected from each individual bat and then stored at −80°C until use.

### RNA extraction, library preparation and sequencing

All samples from each individual bat were homogenized using grinding bowls and rods in MEM medium. The homogenized samples were then centrifuged at 12,000 rpm for 30 min at 4°C to obtain supernatant. Total RNA extraction and purification were performed using the RNeasy Plus universal mini kit (QIAGEN) according to the manufacturer’s instructions. RNA sequencing library construction and ribosomal RNA depletion were performed using the Zymo-Seq RiboFree™ Total RNA Library Kit (Zymo Research). Paired-end (150 bp) sequencing of the 149 dual-indexed libraries was performed on an Illumina NovaSeq platform.

### Virus discovery pipeline

#### Viral genomes assembly and annotation

Raw paired-end sequence reads were first quality controlled, and rRNA reads were removed by mapping against the rRNA database downloaded from the SILVA website (https://www.arb-silva.de/) using Bowtie2. The clean reads were then *de novo* assembled into contigs using MEGAHIT (version 1.2.8)^35^. We performed a blastx search of contigs against the NCBI nr database using Diamond (version 0.9.25)^36^ to roughly classify the sequences by kingdom. The e-value was set at 0.001 to achieve high sensitivity while reducing false positives. Those contigs classified as viruses were used for later analysis. Some viral contigs with unassembled overlaps were merged using SeqMan in the Lasergene software package (version 7.1)^37^. We searched for ORFs in each viral genome using the NCBI ORFfinder (https://ftp.ncbi.nlm.nih.gov/genomes/TOOLS/ORFfinder/), with the genetic code set to standard and with ATG as the only start codon. Then we performed a blastp search against the nr database and manually annotated the viral contigs according to the results.

#### Quantification of virus abundance

We quantified the abundance of each virus in each library as the number of viral reads per million non-rRNA reads (i.e., RPM) by mapping clean non-rRNA reads of each library to the corresponding contigs. To reduce false positives, we applied an abundance threshold of 1 RPM. We further reduced the number of possible false positives from index hopping using the same criterion as described in Shi, et al. ^38^.

#### PCR confirmation of virus genomes

The genome sequence of Bat SARS-like virus BtSY2 was obtained and confirmed by PCR amplification and Sanger sequencing. The WTA product was performed using the Complete Whole Transcriptome Amplification Kit (WTA2)^39^ (Sigma-Aldrich, St. Louis, MO), with the PCR reaction then undertaken using a set of self-designed primer pairs based on the obtained reads. To confirm the recombination breakpoints, long fragments were obtained using the SuperScript IV Reverse Transcriptase and Expand Long Template PCR System.

#### Viral species demarcation and phylogenetic analysis

Viral species were identified based using sequences of the conserved replicase proteins (RNA viruses: RdRp, *Polyomaviridae*: LTAg, *Anelloviridae*: ORF1 protein, *Parvoviridae*: NS1, and other DNA viruses: DNA pol). We applied a 90% cut-off of amino acid sequence similarity to demarcate different virus species. The viruses were aligned using MAFFT (version 7.48)^40^ and ambiguously aligned regions were removed using TrimAl^41^. Phylogenetic trees were then estimated by the maximum likelihood (ML) approach implemented in PhyML version 3.0^42^, employing the LG model of amino acid substitution and the Subtree Pruning and Regrafting (SPR) branch-swapping algorithm. For SARS-related viruses, nucleotide sequences of RdRp, NTD, RBD and N genes were used for phylogenetic analysis, employing the GTR substitution model.

### Recombination analysis of SARS-related viruses

The recombination analysis of SARS-related viruses was performed using similarity plots as implemented in Simplot 3.5.1^43^. The nucleotide sequences of the SARS-related viruses were analyzed with reference strains obtained from GenBank, comprising SARS-CoV Tor2, SARS-CoV-2 Wuhan-Hu-1, as well as some of the closest bat SARSr-CoVs identified so far: Rs4231, WIV16, RaTG13, and BANAL-20-52.

### Protein structure modelling and molecular dynamics (MD) simulations

#### Homology modelling

We built homology models of the Bat SARS-like virus BtSY2 RBD-hACE2 protein complex with MODELLER (version 10.3)^44^, using the known structure of a SARS-CoV-2 RBD-hACE2 complex (PDB ID: 6M0J, resolution 2.45 Å)^45^ as a template. The similarity between Bat SARS-like virus BtSY2 RBD and the template was 97.4%. We removed all NAG and water molecules in the template, and kept the zinc and chloride atoms. We built 100 homology models and selected the top three models based on normalized DOPE score^46^ for the later MD simulations.

#### MD simulation

We used the CHARMM-GUI webservice^47^ to prepare inputs for MD simulations. The three homology models described above and a SARS-CoV-2 RBD-hACE2 complex with known structure (PDB ID: 6M0J) were input to the CHARMM-GUI solution builder pipeline. The four systems were solvated in a water box of 13.5 nm × 9.2 nm × 8.3 nm, with KCl at the concentration of 0.15M. We used CHARMM36m force field^48^ for protein and ions, and TIP3P model^49^ for water.

The models processed by CHARMM-GUI were then used as inputs to GROMACS (version 2022.3)^50^ for MD simulations. The following steps were performed sequentially for each model: (1) energy minimization, (2) 1-ns-long equilibration in NPT ensemble, and (3) 1-ns-long equilibration in NVT ensemble. The temperature and pressure were set to 300K and 1 atm, respectively. We then performed production simulations in NVT ensemble. Production simulation for the top homology model was 1000 ns long, and we performed another two 500-ns-long simulations for the rest two homology models as replications. Similarly, we performed one 1000-ns-long production simulation for the SARS-CoV-2 RBD-hACE2 complex (PDB ID: 6M0J) and two 500-ns-long replications. The settings of these simulations were the same as those described in ref ^51^.

#### Analysis of the MD data

We performed two sets of analyses on the data retrieved from MD. First, we evaluated the stability of RBD-hACE2 binding by measuring deviation of the protein backbones (measured as RMSD) in the duration of simulations, using PLUMED (version 2.7.4)^52^. The backbone RMSD were calculated with respect to energy-minimized structure of each model. We also calculated RMSD separately for RBD, hACE2 and the RBD-hACE2 interface (residues within 0.8 nm to the other subunit in the 6M0J model). Second, we estimated and compared the binding energy of RBD-hACE2 complex using FoldX (version 4)^53^. We visualized the structure of RBD-hACE2 complex using PyMOL (version 2.4.2)^54^.

### Statistical analysis

All statistical analyses were performed in R (version 4.2.0) ^55^

#### Comparing the number of virus species per individual between bat genera

To determine whether specific bat genera harbor more virus species per individual than others, we used a Poisson regression to fit the effect of host genera on number of virus species per individual. We extracted and visualized the estimated effect size of each host genus and its 95% CI. An effect is considered to be significant if its 95% does not include zero.

#### Analysis of cross-species transmission of viruses

To show possible cross-species virus transmission, we visualized the virus-sharing pattern among different bat species using a bipartite network. In this network, a node is either a host or a virus species, and an edge linking a host node and a virus node indicates the presence of that virus in that host. We performed edge betweenness clustering on such network to find network modules, which are subset of nodes such that connections between these nodes are denser than outside of the subset, using the igraph package^56^ in R. A biological interpretation of a network module is that host species within the same module shared more viruses than outside that module.

We performed two sets of statistical tests to further quantitatively evaluate cross-species transmission of viruses among bats. First, we assessed the strength of the correlation of virome composition with both host phylogeny and geographic location using partial Mantel tests implemented in the ecodist package^57^. Differences between virome compositions were represented by Bray-Curtis distance, and phylogenetic distance between hosts was measured as the sum of branch length of pairs of hosts in the COI gene tree. We then used Poisson regression to estimate the effects of (1) phylogenetic distance between hosts, and (2) geographic distance between sample locations on the number of shared virus species between pairs of hosts, including the time intervals between sampling dates to control for its confounding effect.

## DATA AND CODE AVAILABILITY

All meta-transcriptomic sequencing reads have been deposited in SRA database under the project accession (accession id here). The viral genome sequences determined here have been deposited at NCBI GenBank (accession id here). Sample metadata and other materials required to reproduce our results have been deposited together with code and scripts in a GitHub repository (https://github.com/Augustpan/Individual-Bat-Virome). Viral genome sequences are temporarily stored in the above GitHub repository for reviewing, until the NCBI GenBank accessions become available.

## ACKNOWLEDGMENTS

We thank the Alibaba Cloud Computing Co. Ltd. for providing the computational resources for rapid data processing, and the local Centers for Disease Control and Prevention in six trapping sites for their assistance in specimen collection.

MS was supported by Shenzhen Science and Technology Program (KQTD20200820145822023), Guangdong Province “Pearl River Talent Plan” Innovation and Entrepreneurship Team Project (2019ZT08Y464), and Health and Medical Research Fund (COVID190206). YF was supported by the National Natural Science Foundation of China (3156004), Yunnan Reserve Talents for Academic and Technical Leaders of Middle-Aged and Young People (2019HB052), Jianguo Xu Academician Workstation (NO. 2018IC155). ECH was supported by an Australian Research Council Australia Laureate Fellowship (FL170100022) and by InnoHK funding from the Innovation and Technology Commission of Hong Kong. GDL was supported by United States National Institutes of Health U01 AI151810.

## AUTHOR CONTRIBUTIONS

Conceptualization, E.C.H., Y.F., and M.S.; Methodology, J.W., Y.-F.P., W.-H.Y., C.-M.L., W.-C.W., B.L., Y.-L.S., D.-Y.G., G.-D.L., J.L., Y.F., E.C.H., and M.S.; Investigation, J.W., Y.-F.P.,L.-F.Y., W.-H.Y, C.-M.L., J.W., G.-P.K., W.-C.W., Q.-Y.G., G.-Y.X., H.-L.L., Y.-Q.C. E.C.H.,Y.F., and M.S.; Writing – Original Draft, Y.-F.P.; Writing – Review and Editing, J.W., Y.-F.P., L.-F.Y., W.-H.Y, C.-M.L., J.W., G.-P.K., W.-C.W., Q.-Y.G., G.-Y.X., H.-L.L., Y.-Q.C., Y.-L.S., D.-Y.G., Z.-H.G., G.-D.L., J.L., E.C.H. and M.S. Funding Acquisition, J.L., Y.F. and M.S.; Resources (sampling), L.-F.Y., W.-H.Y., Z.-H.G., G.-D.L., and Y.F.; Resources (computational), J.L., E.C.H., and M.S.; Supervision, Z.-H.G., G.-D.L., E.C.H., Y.F., and M.S..

## Supplementary Materials

Supplementary figures S1 ∼ S4

Supplementary tables S1 ∼ S5

