## Supplementary figures S1 ~ S4, Supplementary tables S4 ~ S5 for "Individual bat viromes reveal the co-infection, spillover and emergence risk of potential zoonotic viruses"

**Supplementary Materials**

Supplementary figures S1 ~ S4

Supplementary tables S1 ~ S5

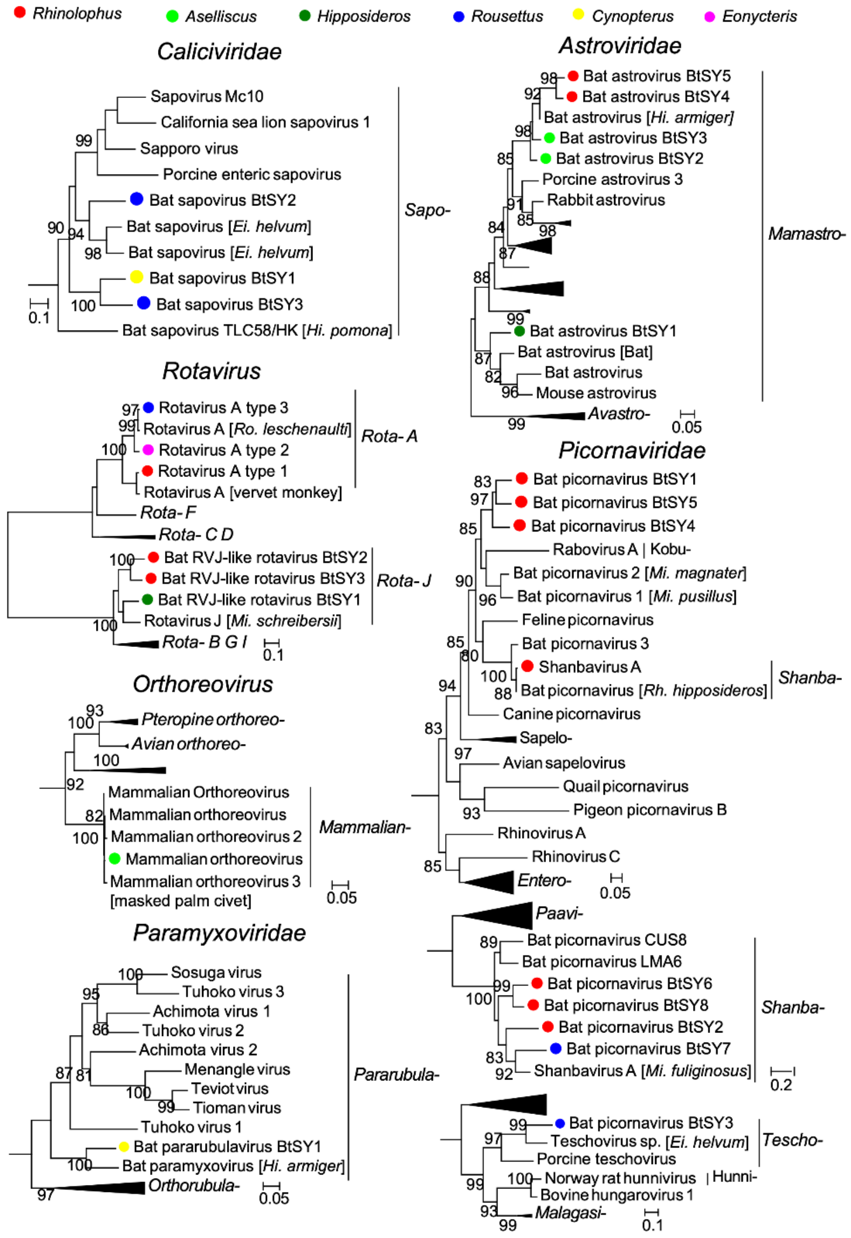


**Fig. S1 |** The evolutionary relationships of the RNA viruses identified in this study. The phylogenetic trees were estimated using a maximum likelihood method based on the RdRp protein. All trees were midpoint-rooted and the branch length indicates number of nucleotide substitutions per site. For clarity, only support values >80% are shown. Dots indicate viruses detected in our samples and colors represent host genus.

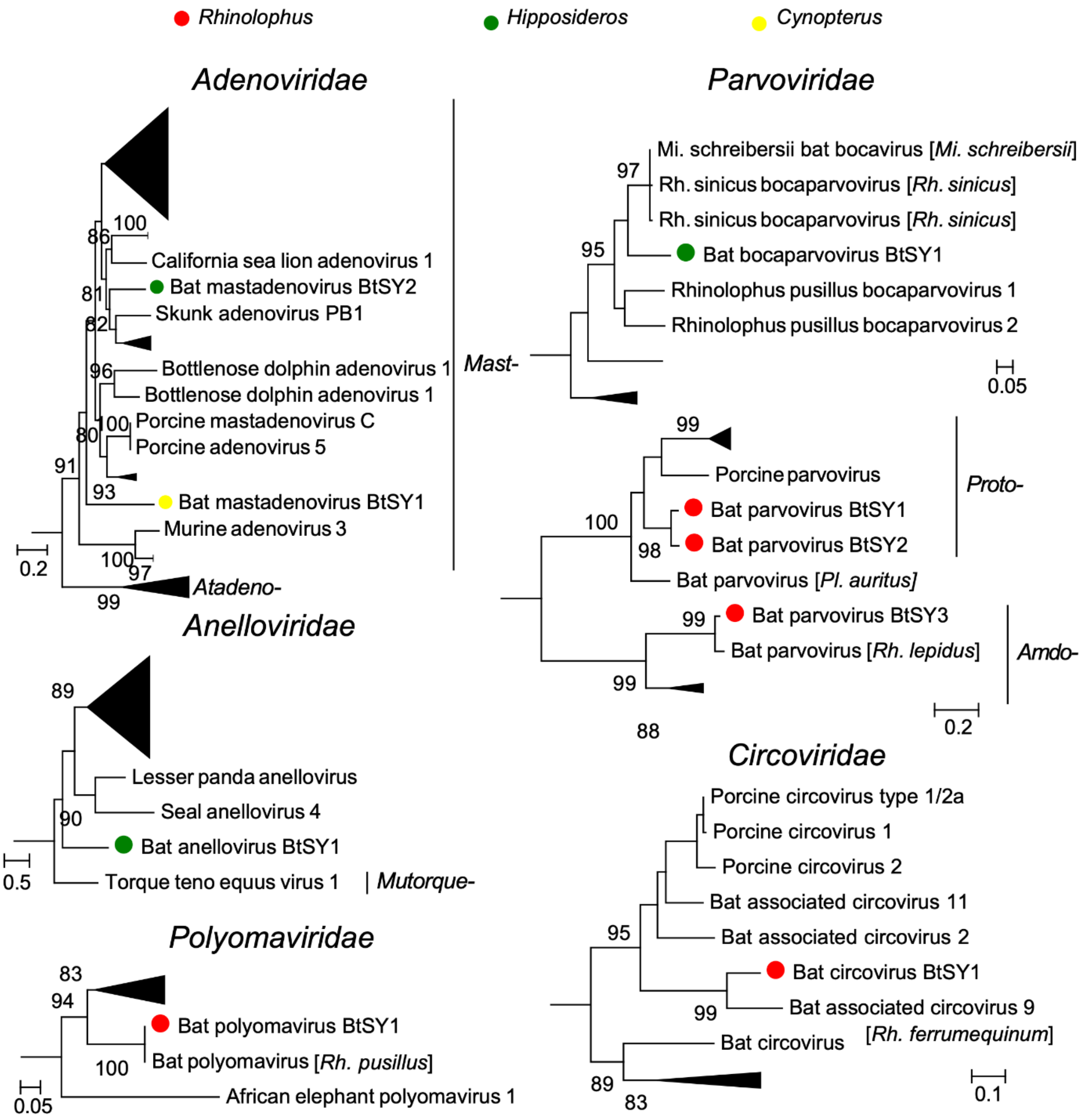


**Fig. S2 |** The evolutionary relationships of the DNA viruses identified in this study. These phylogenetic trees were estimated using a maximum likelihood method based on DNA pol or LTAg (*Polyomaviridae*), ORF1 (*Anelloviridae*), and NS1 (*Parvoviridae*) protein. All trees were midpoint-rooted, and the branch length indicates number of nucleotide substitutions per site. For clarity, only support values >80% were shown. Dots indicate viruses detected in our samples, and colors represent host genus.

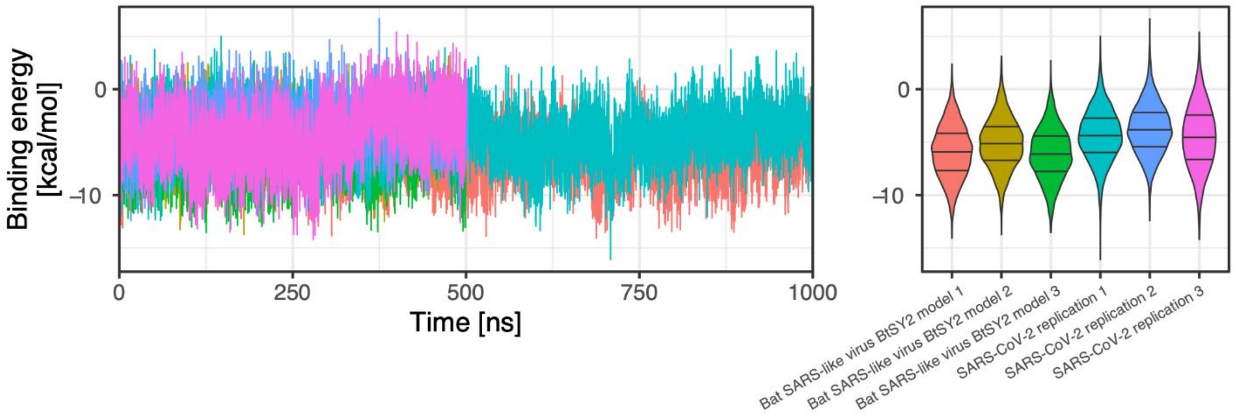


**Fig. S3 |** The predicted binding energy of the RBD-hACE2 complex in the duration of MD simulations. Two 1000-ns-long main simulations and four 500-ns-long replications were performed. These results showed that Bat SARS-like coronavirus BtSY2 RBD can consistently bind to hACE2, and the binding energy is slightly lower than human SARS-CoV-2, suggesting higher affinity.

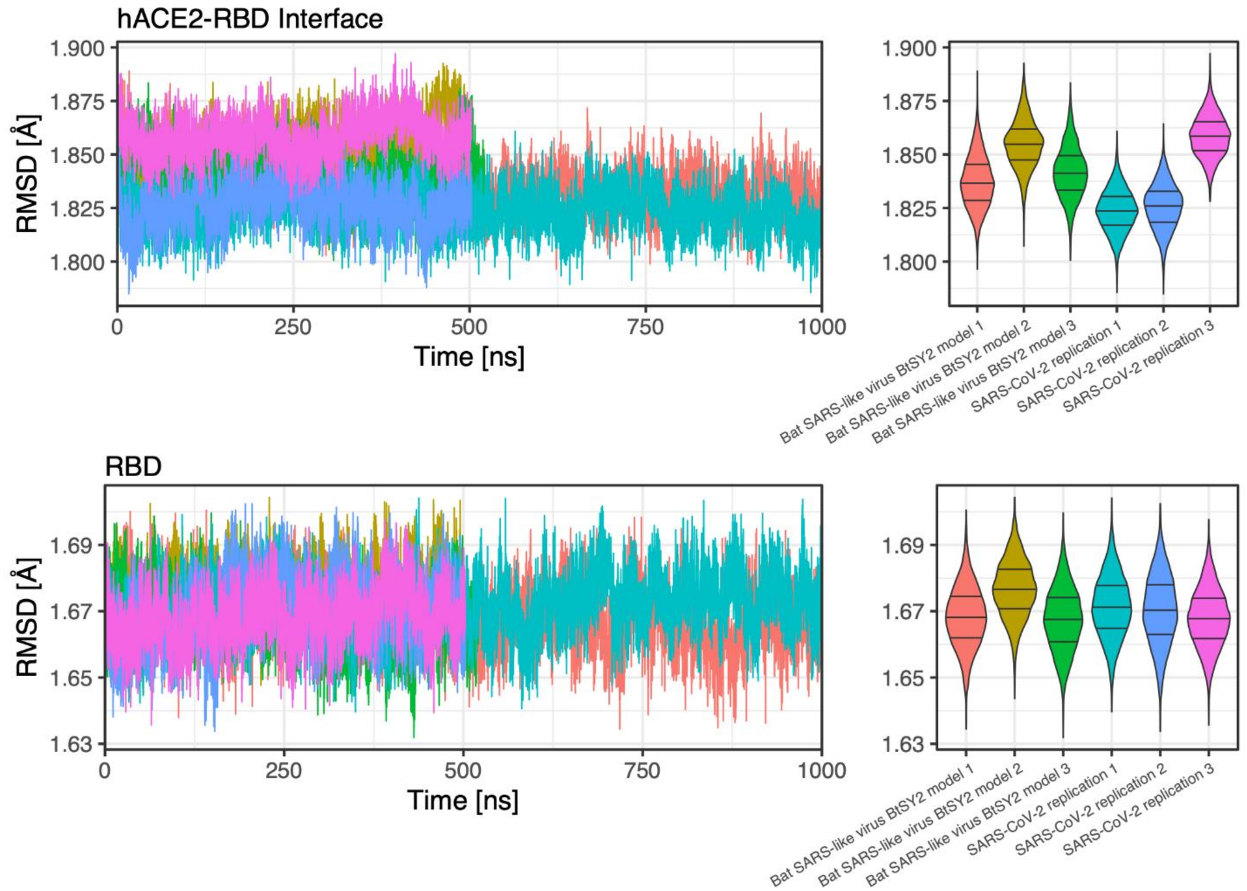


**Fig. S4 |** The deviation in protein backbone position during the MD simulations. RMSD is the abbreviation of root-mean-square deviation of atomic positions. We used backbone (C, N and O atoms in the main chain) RMSD to reflect the stability of RBD-hACE2 binding. Bat SARS-like coronavirus BtSY2 displayed similar binding stability with human SARS-CoV-2, regarding the RBD and RBD-hACE2 interface.

**Supplementary tables**

**Table S1 ~ S3.** Please refer to corresponding spreadsheet files (excel).

**Table S4. Partial Mantel tests showing the effect of host phylogenetic distance, geographic distance and time interval on virome similarity.**

| Variable | Coefficient of correlation | P value |
|---|---|---|
| **Phylogenetic distance** | 0.245 | <0.001 |
| **Geographic distance** | 0.170 | <0.001 |
| **Time interval** | 0.048 | 0.135 |

**Table S5. Identification of five viruses of concern and their prevalence among bats.**

| Virus name | Closest known human or livestock pathogen (amino-acid identity%) | Bat host species | Prevalence (positive/total individuals) |
|---|---|---|---|
| Bat SARS-like virus BtSY1 | SARS-CoV Tor2 (99.6%, RdRp) | *Rh. thomasi* | 2 / 14 |
| | | *Rh. macrotis* | 1 / 2 |
| Bat SARS-like virus BtSY2 | SARS-CoV Tor2 (99.1%, RdRp)<br>SARS-CoV-2 Wuhan-Hu-1 (97.4%, RBD) | *Rh. marshalli* | 1 / 7 |
| | | *Rh. pusillus* | 1 / 16 |
| Rhinolophus bat coronavirus HKU2-like | SADS-CoV GDWT-P83 (99.5%, RdRp) | *Rh. thomasi* | 1 / 14 |
| | | *Rh. pusillus* | 1 / 16 |
| Mammalian orthoreovirus | Porcine reovirus SHR-A (99.1%, RdRp) | *As. stoliczkanus* | 3 / 35 |
| | | *Hi. armiger* | 3 / 11 |
| | | *Hi. larvatus* | 1 / 13 |
| | | *Rh. marcrotis* | 1 / 2 |
| Rotavirus A type 1 | Human rotavirus AU-1 (98.1%, RdRp) | *As. stoliczkanus* | 9 / 35 |
| | | *Rh. thomasi* | 1 / 14 |
| | | *Rh. pusillus* | 1 / 16 |
| | | *Rh. marshalli* | 1 / 7 |
| | | *Rh. pearsonii* | 1 / 2 |
